# Hybrid Support Vector Regression Model and K-Fold Cross Validation for Water Quality Index Prediction in Langat River, Malaysia

**DOI:** 10.1101/2021.02.15.431242

**Authors:** Naeimah Mamat, Firdaus Mohamad Hamzah, Othman Jaafar

## Abstract

Water quality analysis is an important step in water resources management and needs to be managed efficiently to control any pollution that may affect the ecosystem and to ensure the environmental standards are being met. The development of water quality prediction model is an important step towards better water quality management of rivers. The objective of this work is to utilize a hybrid of Support Vector Regression (SVR) modelling and K-fold cross-validation as a tool for WQI prediction. According to Department of Environment (DOE) Malaysia, a standard Water Quality Index (WQI) is a function of six water quality parameters, namely Ammoniacal Nitrogen (AN), Biochemical Oxygen Demand (BOD), Chemical Oxygen Demand (COD), Dissolved Oxygen (DO), pH, and Suspended Solids (SS). In this research, Support Vector Regression (SVR) model is combined with K-fold Cross Validation (CV) method to predict WQI in Langat River, Kajang. Two monitoring stations i.e., L15 and L04 have been monitored monthly for ten years as a case study. A series of results were produced to select the final model namely Kernel Function performance, Hyperparameter Kernel value, K-fold CV value and sets of prediction model value, considering all of them undergone training and testing phases. It is found that SVR model i.e., Nu-RBF combined with K-fold CV i.e., 5-fold has successfully predicted WQI with efficient cost and timely manner. As a conclusion, SVR model and K-fold CV method are very powerful tools in statistical analysis and can be used not limited in water quality application only but in any engineering application.

## Introduction

Water pollution is the contamination of water bodies such as lakes, rivers, oceans, and groundwater. This problem is highly attributed by water pollutants which are discharged directly or indirectly into the water bodies without proper treatment. As a result, a lot of efforts have been done by researchers to develop prediction models for water quality parameters. Among the modelling methods used are statistical modelling such as multivariate analysis [4,6,18,20,23] and Artificial Intelligent (AI) modelling [15]. As time passes, researchers developed Machine learning (ML) tools base on AI architecture. ML is a subfield of the AI which deals with the methods development for allowing the computer processor to calibrate the natural behaviour and learn its nonlinearity process [3]. Several techniques were developed for modelling engineering applications and natural behaviour utilizing ML such as Artificial Neural Networks (ANN) [10,12,13,17,26], adaptive neuro-fuzzy system (ANFIS) [50-52,55], Decision Tree [12,17], Ensemble methods [25], K-means [27] and Support vector machines (SVM) [1,2,3,24,25,27,29]. Despite having complex calculation, nonlinear and stochastic in nature, ML models can be used to predict water quality effectively in addressing the difficulty of monitoring water quality parameters [17].

Water surface pollution has been identified as one of major problem in Sungai Langat. The increasing in human population in this area causes the increasing in waste to the drainage system. Hence, the unwanted waste will be flushed directly into the river. Department of Environment (DOE), under the Ministry of Natural Resources and Environment has undertaken various efforts and programs to continuously monitor water quality levels. Based on water quality data, the water quality index (WQI) is calculated to assess the status and the class of river water quality. Several factors need to be considered to predict the degree of change in water quality. The WQI calculation includes six water quality variables, namely SS, pH, AN, dissolved oxygen (DO), chemical oxygen demand (COD), and biochemical oxygen demand (BOD) [35]. Therefore, the development of water quality prediction model is a very important task towards better river water quality management. By considering the importance of environmental preservation, this research mainly focused on the development a model for predicting the class of WQI which indicate the status of river water quality. In this work, the main objective is to utilize a hybrid of Support Vector Regression (SVR) modelling and K-fold cross-validation as a tool for WQI prediction. Base on the literature review, the studies comparing SVM methods with Decision Tree and ANN showed that SVM models provide better predictive accuracy [16,49].

### Area study and data set

The map in Fig 1 shows the location of Langat River located in the state of Selangor, Malaysia. The Langat River basin has a total catchment area of approximately 1815 km^2^ and lies within longitude 101°17′E to 101°55′E and latitudes 2°40′N to 3°17′N. The Langat River basin is one of the most important basins which supply water to about a third of the state of Selangor. The Langat River flows from the Titiwangsa Range in the Northeast of Hulu Langat District and drains into the Straits of Malacca. Referring to Department of Environment [21], there are eight Langat sub-basin, namely Teluk Panglima Garang, Teluk Datok, Putrajaya, Kajang, Cheras, Hulu Langat, Pangsoon and Ulu Lui. In this study, the largest sub basin, Kajang was chosen. The sub basin lies in the middle section of Langat River, with fluctuating water quality. Hafizan Juahir et al. [14] states that the downstream section of Langat River has been identified among 42 tributaries in Peninsular Malaysia that have been categorized as very polluted. Hence, the water quality is a great concern since river water is an important source for the citizen in Langat River.

**Fig 1:**
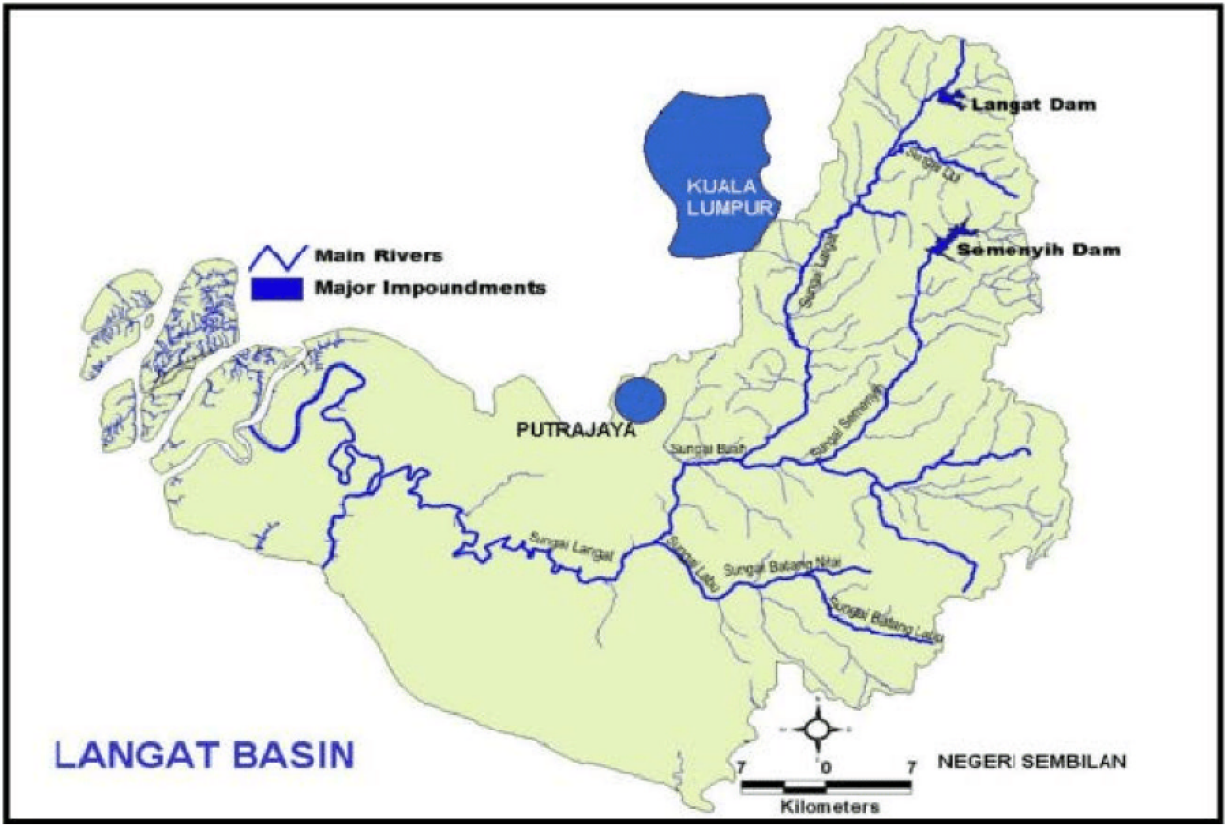
Map of the Langat River basin located in Selangor, Malaysia.

In this study, two water quality stations in Langat river, Kajang have been selected which are station 1L15 and 1L04 act as upstream and downstream point, respectively. Location of the sampling stations are illustrated in Fig 2 while the coordinates of the stations are shown in Table 1.

**Fig 2:**
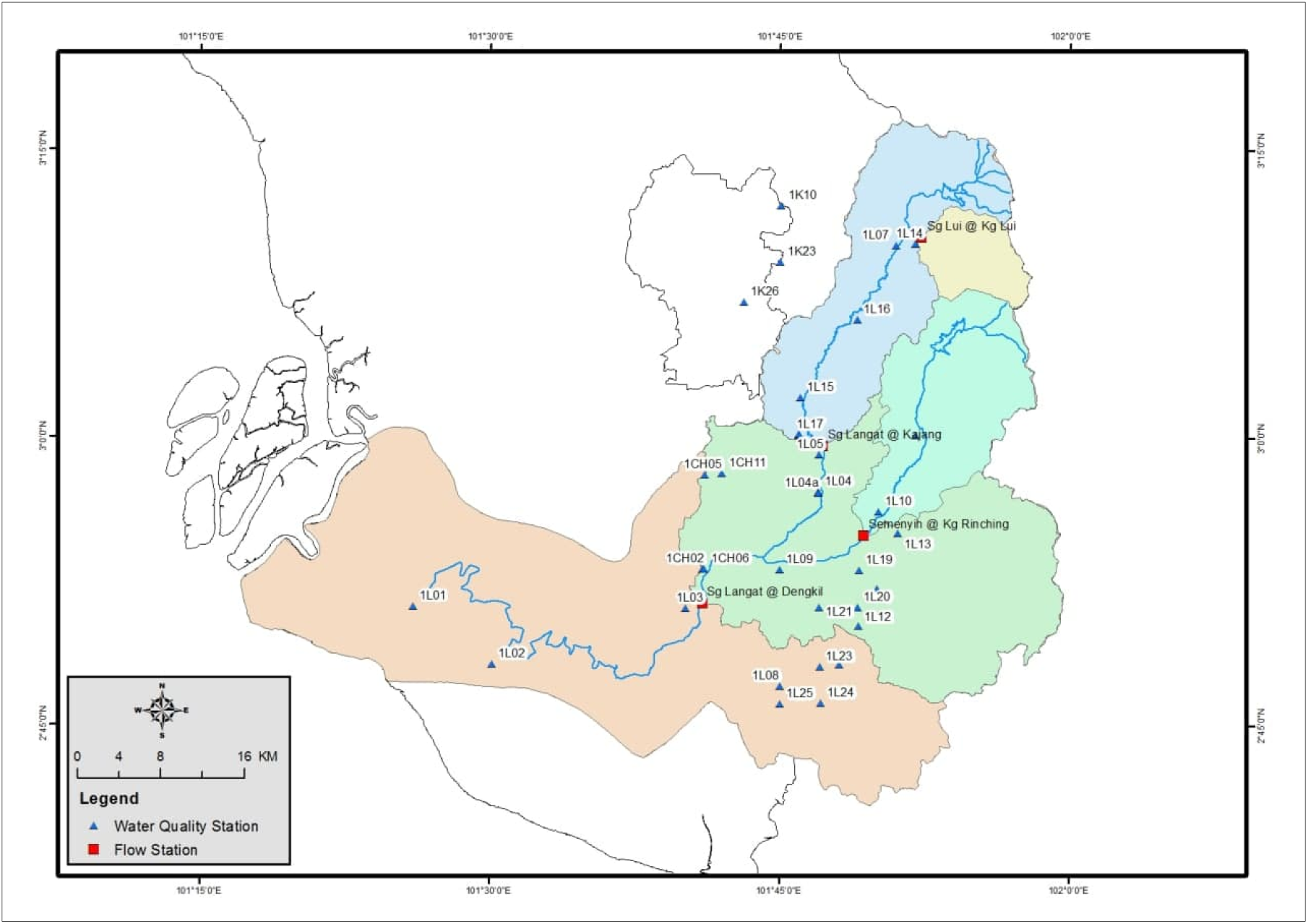
Map of the water quality monitoring stations.

**Table 1:**
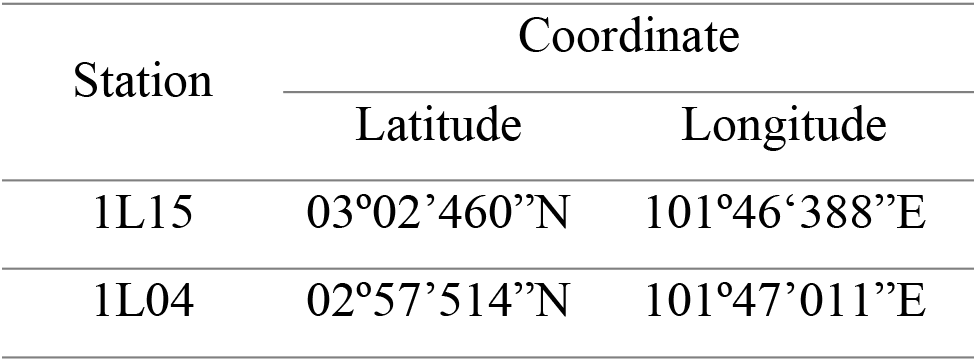
Coordinates of sampling station

### Methodology

#### Water Quality Index (WQI)

Water Quality Index (WQI) has been used to check the status of river quality for different uses. WQI was developed by Brown et.al 1970 and then, in 1975, was improved by the National Sanitation Foundation (NSF). WQI is a mathematical formula that is a combination of several parameters and presents in one number [41]. Generally, WQI is a number with no units and in the range of 0 and 100. Higher index value represents better water quality [47].

The Department of Environment (DOE) is a government agency responsible for monitoring the condition of rivers in Malaysia and developing water quality guidelines along with the identification of pollution sources. The quality level of water in the Malaysian river is calculated based on the Malaysian Department of Natural Water Quality Index (DOE-WQI) and is classified according to the Malaysian Interim State Water Quality Standard (INWQS). DOE-WQI is a river water assessment related to pollution load categorization while INWQS provides classifications according to useful uses. The implementation of DOE WQI has been practiced in Malaysia for over 30 years and the calculation of WQI is based on the opinion-poll formula [43].

WQI is calculated based on dissolved oxygen (DO), biochemical oxygen demand (BOD), chemical oxygen demand (COD), ammoniacal nitrogen (AN), suspended solid (SS) and pH. Among the six parameters, DO carries the highest weightage of 0.22 and pH carries the lowest of 0.12 in WQI equation [48]. The calculation is using the following equation:

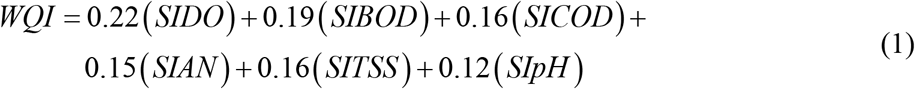

The parameter value is converted into sub-index (SI) before calculating WQI and Table 2 showing the sub-index calculation of the WQI parameter. The guidelines set by the DOE which is a river with WQI values in the 0-59%, 60-80%, and 81-100% considered to be polluted, slightly polluted, and clean, respectively [48]. There are five classification of water quality based on WQI value. Class ranges and class definitions are shown in Table 3 and Table 4, respectively.

**Table 2:**
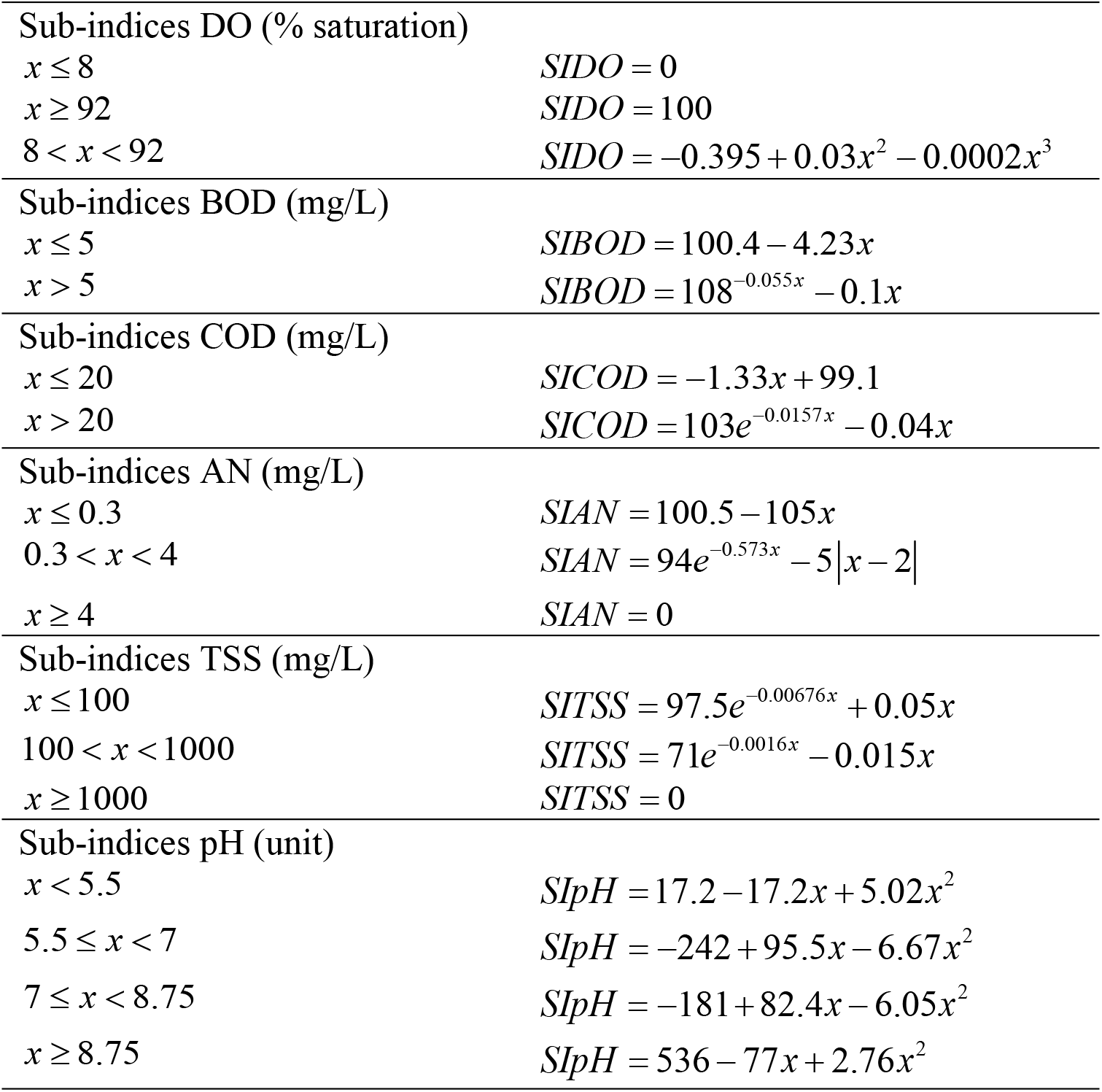
Calculation of WQI [48]

**Table 3:** INWQS classes [48]

| Parameter | Unit | Classes |  |  |  |  |
| --- | --- | --- | --- | --- | --- | --- |
|  |  | I | II | III | IV | V |
| Ammoniacal Nitrogen (AN) | mg/l | <0.1 | 0.1-0.3 | 0.3-0.9 | 0.9-2.7 | >2.7 |
| Biochemical Oxygen Demand (BOD) | mg/l | <1 | 1-3 | 3-6 | 6-12 | >12 |
| Chemical Oxygen Demand (COD) | mg/l | <10 | 10-25 | 25-50 | 50-100 | >100 |
| Dissolved Oxygen (DO) | mg/l | >7 | 5-7 | 3-5 | 1-3 | <1 |
| pH | unit | >7.0 | 6.0-7.0 | 5.0-6.0 | <5.0 | >5.0 |
| Total Suspended Solids (TSS) | mg/l | <25 | 25-50 | 50-150 | 150-300 | >300 |
| Water Quality Index (WQI) |  | >92.7 | 76.5-92.7 | 51.9-76.5 | 31.0-51.9 | <31.0 |

**Table 4:** Definition of Classes for INWQS [48]

| Class | Uses |
| --- | --- |
| Class I | Conservation of natural environment<br>Water Supply I – practically no treatment necessary (except by disinfection or boiling only)<br>Fishery 1 – very sensitive aquatic species<br>Water Supply II – conventional treatment required. |
| Class IIA | Fishery II – sensitive aquatic species<br>Recreational use with body contact |
| Class IIB | Water Supply III – extensive treatment required. |
| Class III | Fishery III – common, of economic value, and tolerant species; livestock drinking. |
| Class IV | Irrigation |
| Class V | None of the above |

#### Water Quality Index (WQI) Parameters

The water-quality data utilized in this research is obtained through the Malaysian Department of Environment (DOE). Ten years monthly time-series data for selected parameter sets through two monitoring stations are utilized in this research. Six water-quality parameters are selected for SVM modelling in this research, specifically pH, Suspended Solids (SS), Dissolved Oxygen (DO), Ammonia Nitrogen (AN), Chemical Oxygen Demand (COD), and Biochemical Oxygen Demand (BOD).

Dissolved oxygen concentration is used as main indicator for the health of a river or water body. It represents the ability of the water body to support plant and animal life. In the other hands, BOD refers to a measure of total amount of oxygen removed from water biologically or chemically in a specified time and at a specific temperature. BOD indicates the total concentration of DO utilised either during degradation of organic matter or the oxidation of inorganic matter. In addition, COD is a measure of the amount of oxygen reduction due to chemical oxidation of pollutants in the system: it involves the addition of a chemical oxidising agent such as potassium permanganate or dichromate to a water sample for a standard period [54].

Organic nitrogen represents nitrogen which is present in organic matter in the form of compounds such as proteins and amino acids. These compounds are hydrolysed by bacteria to form ammonium compounds. As with BOD, the organic nitrogen hydrolyses at a fast and a slow rate represented by temperature-dependent, first-order kinetics. Ammoniacal nitrogen represents nitrogen which exists in the form of ammonia or ammonium ions. It can be formed by the hydrolysis of organic nitrogen, as described above, but also enters the river system directly from industrial or sewage effluent. Ammoniacal nitrogen is oxidised to nitrite by Nitrosomonas bacteria [54].

Total suspended solids (TSS) is usually referred to the particles in water which is usually larger than 0.45 μm. Many pollutants can be attached to TSS, which is not good for the aquatic habitat and lives. High suspended solids also prevent sunlight to penetrate water. Total dissolved solid (TDS) consists of dissolved minerals and indicates the presence of dissolved materials that cannot be removed by conventional filtration. The presence of synthetic organic chemicals imparts offensive tastes, odours and colours to fish and aquatic plants even when they are present in low concentrations [47]. As for pH, this parameter directly measures the activity of the hydrogen ion, H+. The lower the pH, the higher the H+ activity and the more acidic is the water [53]. The neutral pH is considered as 7.0 [48]. Descriptive statistics for measured water-quality parameters in the Langat River from are displayed in Table 5 below.

**Table 5:** Descriptive statistics of the selected water quality parameter in Langat River.

| Station | Basic Statistics | DO (mg/L) | BOD (mg/L) | COD (mg/L) | SS (mg/L) | pH (unit) | AN (mg/L) | WQI |
| --- | --- | --- | --- | --- | --- | --- | --- | --- |
| L15 | Mean | 6.63 | 6.02 | 35.26 | 400.04 | 7.05 | 1.69 | 69 |
|  | Min | 2.71 | 1.00 | 5.00 | 48.00 | 5.43 | 0.01 | 45 |
|  | Max | 8.67 | 37.00 | 114.00 | 2420.00 | 7.92 | 7.52 | 87 |
|  | SD | 0.99 | 5.68 | 17.89 | 481.09 | 0.38 | 1.39 | 8.50 |
|  | CV | 0.15 | 0.94 | 0.51 | 1.20 | 0.05 | 0.82 | 0.12 |
| L04 | Mean | 5.09 | 7.33 | 38.12 | 207.35 | 7.10 | 2.00 | 64 |
|  | Min | 0.15 | 2.00 | 11.00 | 14.00 | 5.85 | 0.01 | 27 |
|  | Max | 8.79 | 37.00 | 159.00 | 1190.00 | 7.83 | 6.23 | 83 |
|  | SD | 1.50 | 5.03 | 20.34 | 198.13 | 0.34 | 1.44 | 10.94 |
|  | CV | 0.29 | 0.69 | 0.53 | 0.96 | 0.05 | 0.72 | 0.17 |

#### Structure of Regression in Support Vector Machines (SVM)

Support vector machines have been introduced by Boser et.al [4] and been developed to solve the problem of classification and been extended to the regression problem [15,26]. Support vectors are the training points that are the nearest to the separating hyperplane and the basic concept of SVM is illustrated in Fig 3. There are decision functions that are accountable for example, hyperplanes that can delineate the positive and negative data that has marked the maximum margins. This shows the range from the nearest positive sample to a hyperplane and the range between the nearest negative sample and the hyperplane shall be maximized [1-2]

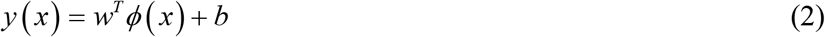

where, *ϕ*(*x*) represents the high dimensional feature spaces, which is nonlinearly mapped from the input space *x*. The coefficients are *w* and *b* are estimated by minimizing the regularized function *R*(*C*):

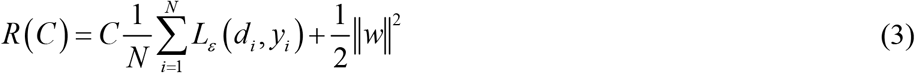

where

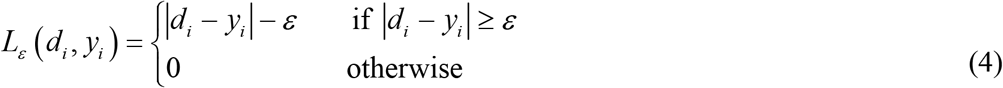

**Fig 3:**
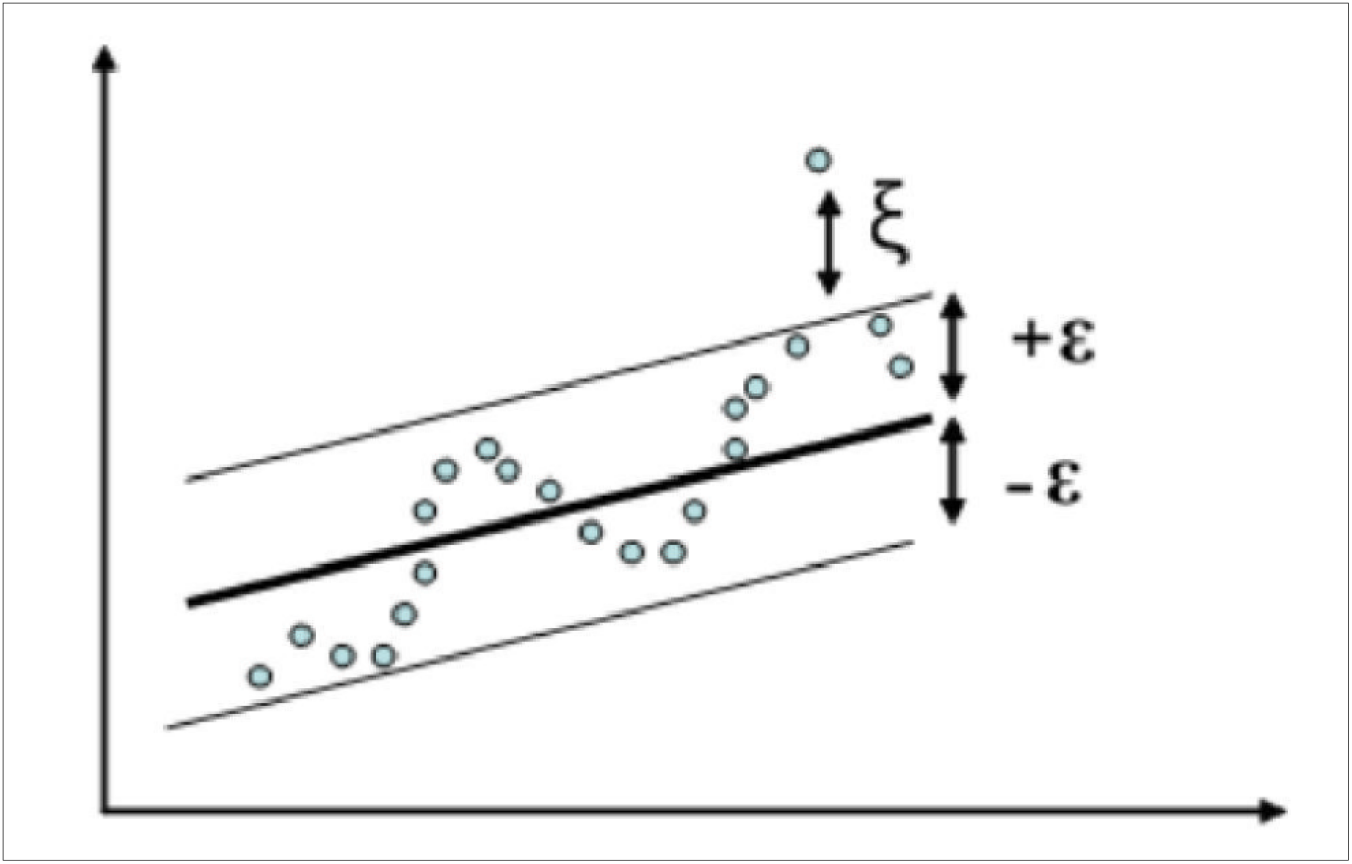
Hyperplane and the basic concept of support vector machine (SVM) [1-2].

To obtain the estimation of *w* and *b*, Equation (3) is transformed to the primal function given by Equation (5) by introducing the positive slack variables *ξ_i_* and 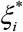 as follows:

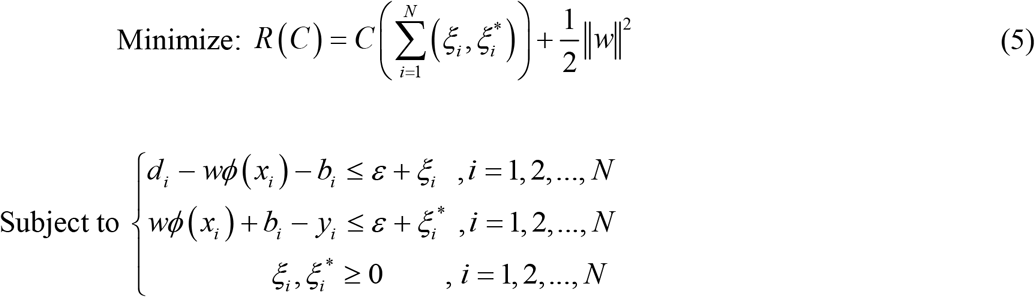

The first term 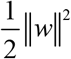 is the weights vector norm, *d_i_* the desired value, and *C* is referred as regularized constant determining the trade-off between the empirical error and the regularized term. *ε* is called the tube size of SVM as shown in Fig 3.

It is similar to the equation accuracy that is related in the training data. The multipliers that are non-zero are known as support vectors. This is where the variable slacks and brought into the study. *ξ_i_* and 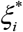 are introduced and by introducing this and by exploiting constraint optimality the function decision by Equation (1) gives the following:

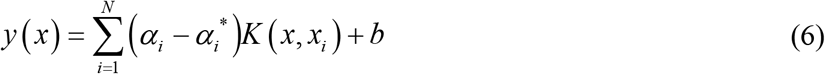

In Equation (6), *α_i_* and 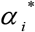, are the so-called Lagrange multipliers. They satisfy the equalities 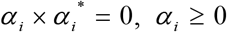 and 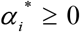 where, *i* = 1,2,…, *N*, and are obtained by maximizing the binary function of Equation (5) which has the following form:

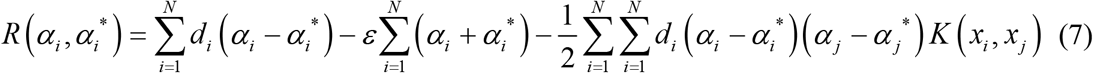

with the constraints

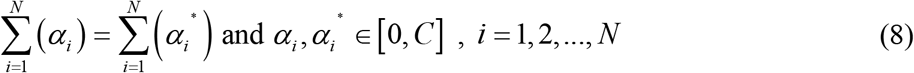

*K* (*x_i_*, *x_j_*) is defined as the kernel function. The value of the kernel is equal to the inner product of two vectors *x_i_* and *x_j_* in the feature space *ϕ*(*x_i_*)and *ϕ*(*x_j_*), that is, *K*(*x_i_,x_j_*) = *ϕ*(*x_i_*)×*ϕ*(*x_j_*).

Four common kernel function types of SVM are given as follows:

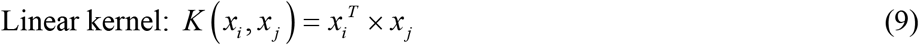

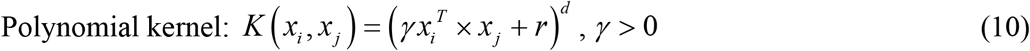

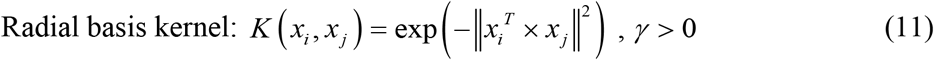

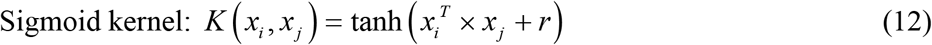

Here *γ, r* and *d* are kernel parameters. The kernel parameters should be carefully chosen as it indirectly defines the structure of the high dimensional feature space *ϕ*(*x*) and thus controls the complexity of the final solution [1].

There are two types of SVM regression; both have the general formula given in Equation (2). The first type of SVM regression is known as Type 1 or *Epsilon.* This type of error function is given by the formula shown in Equation (6). The second type of regression is known as *Nu.* C and gamma are the parameters in the RBF kernel for a nonlinear SVM. As for a higher value of gamma, the variance is smaller indicating the support vector does not have wide-spread influence. In general, larger gamma leads to higher bias and lower variance models, and vice-versa.

Meanwhile, C is the parameter for the soft margin cost function, which controls the influence of each individual support vector, whereas this process involves trading error penaltyfor stability. The higher C will cause the more SVM adjusting to the training set and finally will face the risk of overfitting. Technically, higher value of C leads to lower bias and higher variance models, and smaller C will cause higher bias and lower variance [5]. Fig 4 below shows the flow of algorithm of SVM as explained earlier.

**Fig 4:**
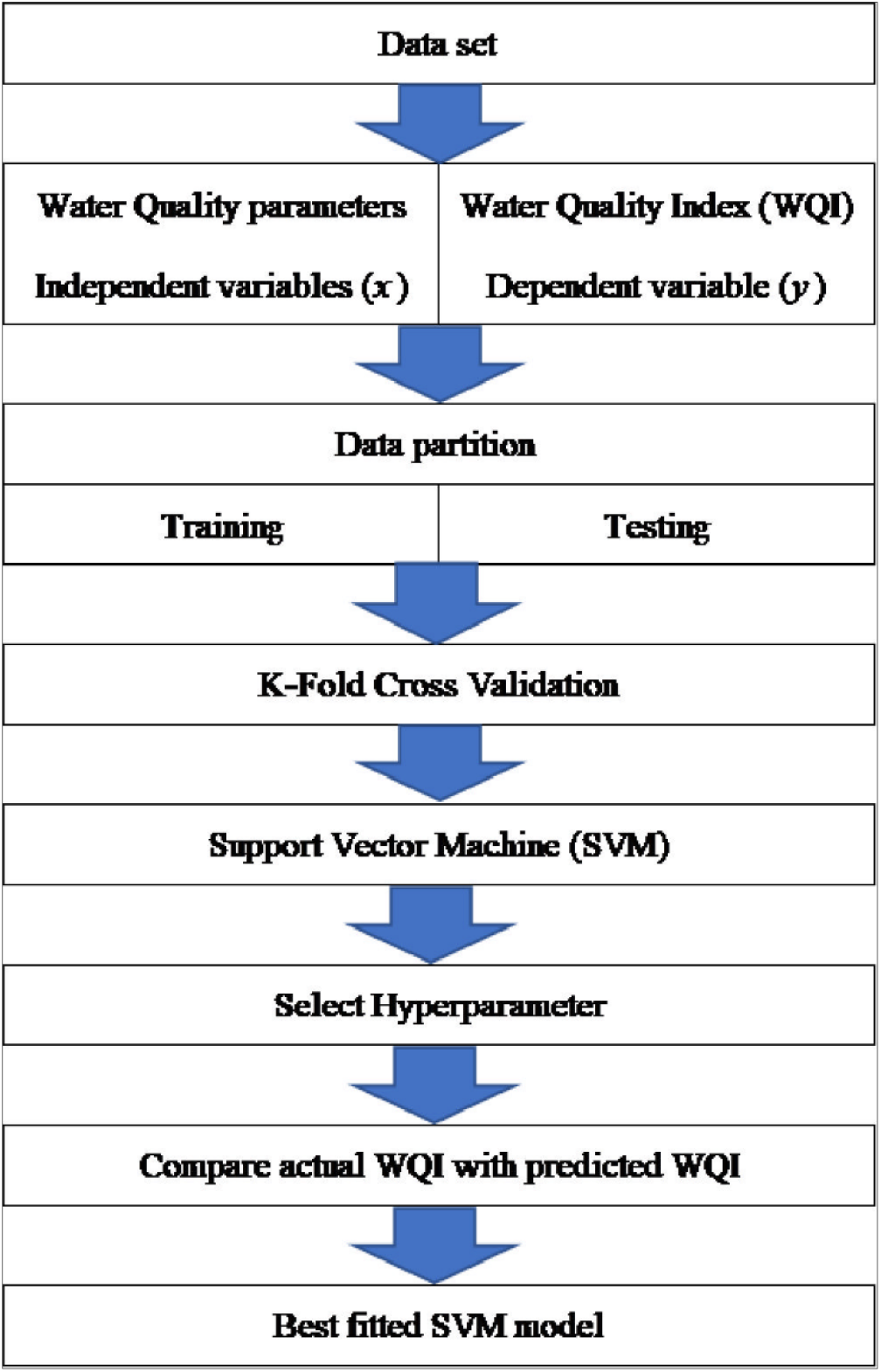
The flow of algorithm of SVM model.

#### Statistical Performance of SVR model

Performance of SVR models were evaluated based on these criteria of indicators [2,36-37].

1. Nash Sutcliffe efficiency (NSE) or Coefficient of Efficiency (CE) To evaluate model performance, Nash Sutcliffe efficiency (NSE) was utilized.

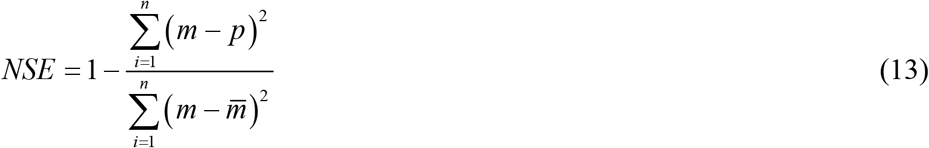

where *n* is the number of observations, *m* and *p* correspond to the measured and predicted data, while 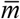 denotes the average of the measured data.
2. RMSE, Root Mean Square Error (MSE) Root Mean square error (RMSE) is utilized to establish and define the fit of the network outputs to the desired scheme. Lower values for RMSE enable higher performance (values close to 0 indicate reliable and accurate results), which is expressed as follows:

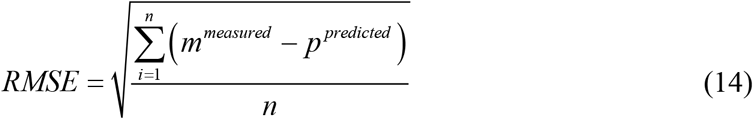
3. Pearson correlation coefficient (R) Pearson correlation coefficient (*R*) is utilized to measure the strength and direction of the linear relationship between the observed and predicted data. Values can range from −1 to +1. It is defined as follows:

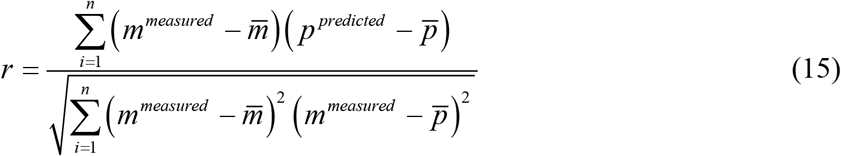
4. MAE The mean absolute error is the average difference between imputed and observed data points. Mean absolute error (MAE) ranges from 0 to infinity and a perfect fit is obtained when MAE equals to 0. MAE is the average absolute difference between two variables designated *m_i_* and *p_i_*. MAE is fundamentally easier to understand.

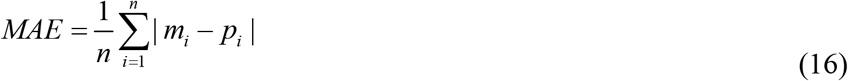

#### K-fold cross validation of SVR model

Cross validation (CV) is one of the techniques that widely used to evaluate and to compare machine learning algorithms such as SVM model. CV is also known as a resampling method because it involves fitting the same statistical method multiple times using different subsets of the data. In cross validation, the data set will be partitioned into two sections which are the first part is for training the model and the second part is for testing the model. An accuracy of the model will be determined by estimating the prediction error. One of the approaches in cross validation is k-fold cross validation.

The k-fold cross-validation evaluates the SVM model performance on different subset of the training data and then compute the average prediction error rate. The algorithm (Fig 5) is started by splitting the data randomly into k number of folds. Afterwards, k iteration of training and testing are performed and then the preferred type of SVM model is presented in sequence which occurs to the k-1 fold [24]. In the first iteration, the first fold is used to test the model and the rest are used to train the model. In the second iteration, the second fold is used for testing set while the rest as the training set. This process is repeated until each fold of the k-fold has been used as the testing set. After building the model in a training phase, the model will be applied on a new unseen observations or test data set which is not used to build the model. Then the prediction error will be computed.

**Fig 5:**
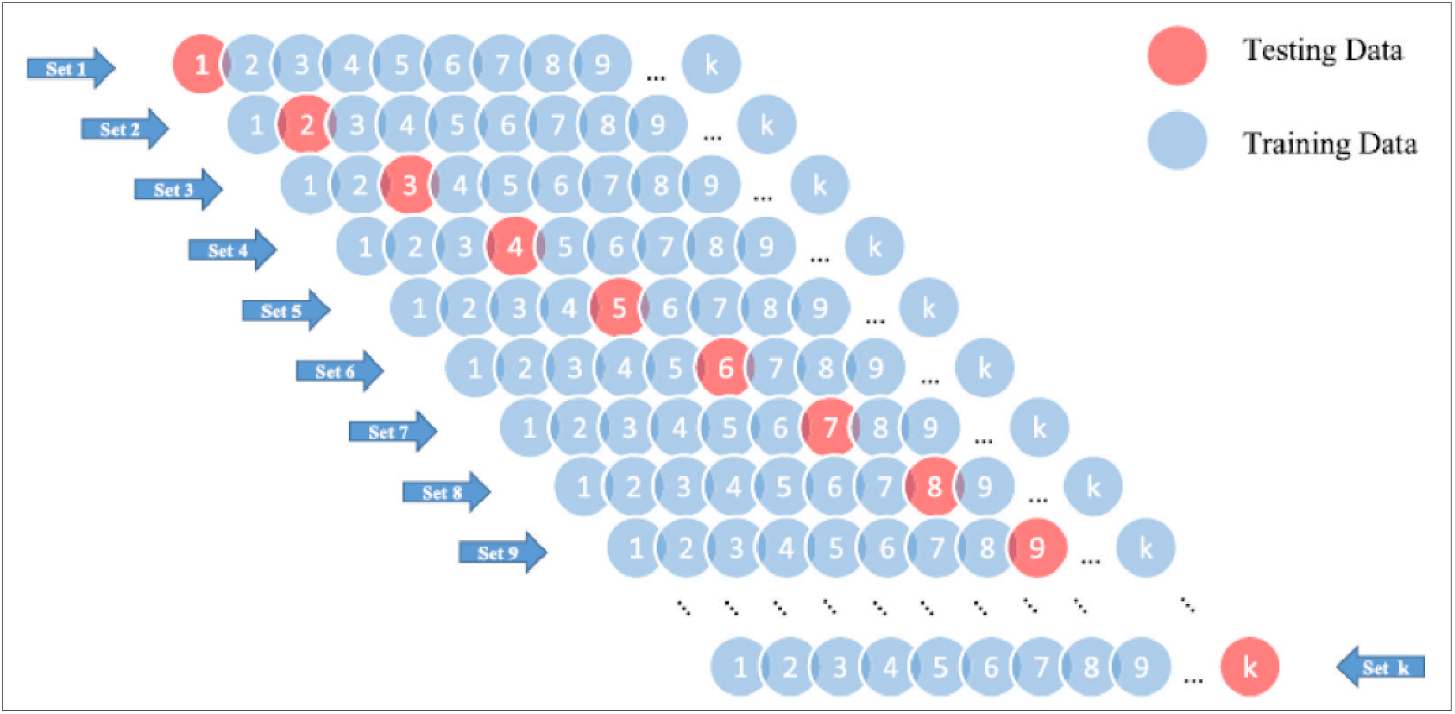
Schematic diagram of K-Fold cross validation procedure [10].

K-fold cross-validation (CV) is a robust technique to evaluate the accuracy of a model. The advantage of k-fold CV is always gives more accurate estimates of the test error rate [21]. Smaller value of K is more biased and therefore unacceptable. Instead, larger value of K is less biased, but can affected to higher variance. Even though, there is no formal rule to choose the value of k, but by considering this situation, the common choices of k are 5 [37] or 10 (Fig 6). These values have been shown empirically to yield test error rate estimates that suffer neither from excessively high bias nor from very high variance. [12,18,19].

**Fig 6:**
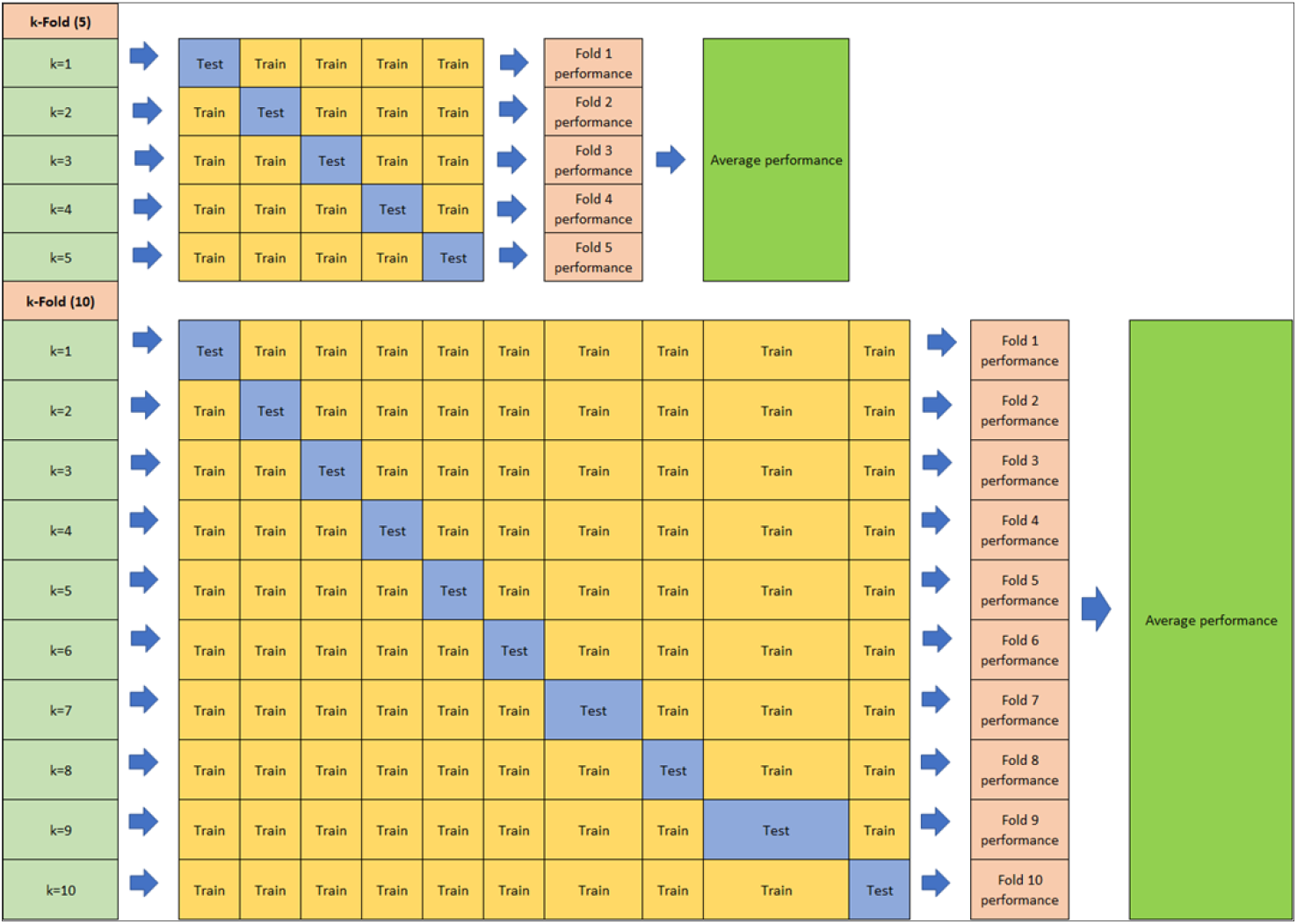
Schematic diagram of 5-Fold cross validation and 10-Fold cross validation procedure.

## Results and discussion

In this study, the SVM model for the prediction of WQI has been developed by using various of kernel functions such as Linear, Radial Basis, Polynomial and Sigmoid function. Initially, these functions need to be evaluated to determine the best kernel type by utilizing 10-fold cross validation value. As a result, a comparison for the performance of the SVM models has been shown in Table 2. In the table, RBF kernel function shows the best value of correlation coefficient in the testing data session (0.987) then followed by Linear (0.942), Polynomial (0.896) and Sigmoid kernel function (−0.031). Therefore, the RBF function will be utilized for further analysis.

**Table 2:**
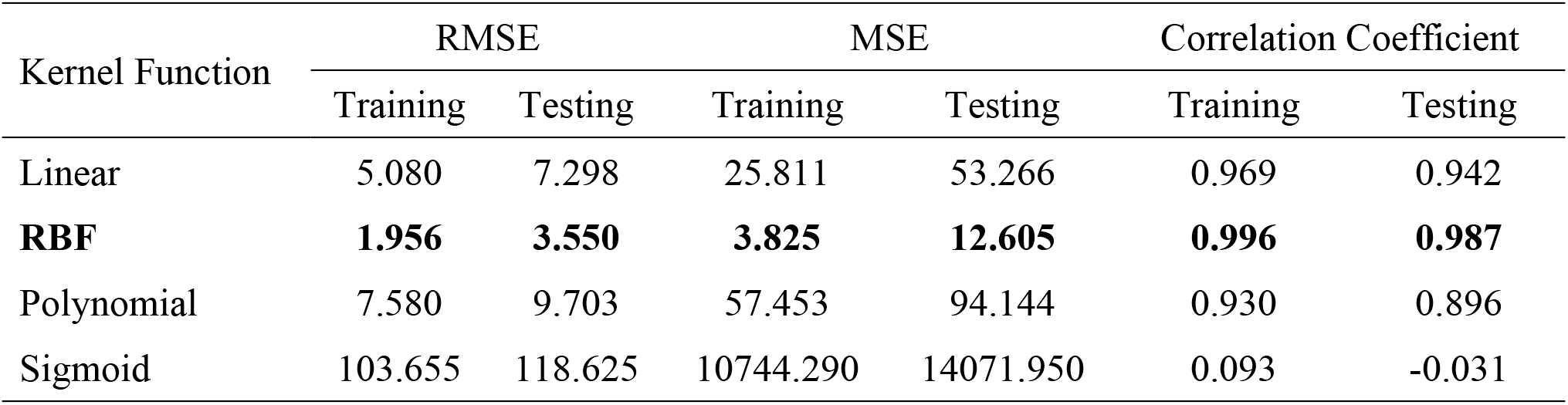
Performance of SVM model with four kernel functions.

The selection of the optimal parameter sets is an important role in order to attain valid predictive performance for SVM model. The SVM generalization performances are influenced by setting of hyper-parameters *(C, γ)* and kernel parameters (epsilon ε, nu v). Hence, in order to find the best SVM performance, two types of RBF kernel functions which are epsilon ε and nu v are considered for WQI prediction. As for epsilon-RBF model, *C* is fixed to 8 and gamma equals to 0.2 and epsilon is set to various values in the range of 0.001 to 0.5. As a result, the training and the testing phases shows as epsilon increases, the value of RMSE also increases. However, the value of correlation coefficient and the number of support vectors decreases. Table 3 below shows all the related result. Consequently, epsilon equals to 0.001 is selected for further analysis. This is because the value shows the minimum generalization error with acceptable number of support vectors.

**Table 3:** Result of various epsilon parameter, ε of SVM model where C=8 and gamma=0.2 of total behaviour prediction model.

| Epsilon<br>$\epsilon$ | RMSE | | Correlation Coefficient | | | No. of Support<br>Vectors |
| --- | --- | --- | --- | --- | --- | --- |
|  | Training | Testing | Training | Testing | Overall |  |
| **0.001** | **0.3233** | **1.9869** | **0.9993** | **0.9752** | **0.9936** | **83** |
| 0.1 | 0.7773 | 3.2411 | 0.9966 | 0.9327 | 0.9820 | 56 |
| 0.2 | 1.4004 | 2.7540 | 0.9885 | 0.9543 | 0.9803 | 42 |
| 0.3 | 1.9332 | 2.9198 | 0.9781 | 0.9518 | 0.9710 | 29 |
| 0.4 | 2.4859 | 3.3201 | 0.9630 | 0.9305 | 0.9547 | 25 |
| 0.5 | 3.0320 | 3.9170 | 0.9425 | 0.8868 | 0.9301 | 22 |

As for nu-RBF model, gamma is fixed to 0.2, *C* equals to 8 and the nu in the range of 0.001 to 0.5. These fixed values then integrated in the algorithm and as a result, the accuracy of the performance for the nu-RBF model is shown in Table 4 below. In the table, as nu value increases, the predictive accuracy of the test data slightly increases along with the correlation coefficient and the number of support vectors. When the value of nu reached the point 0.3 and 0.4, the RMSE of the training data remain unchanged. Thus, nu=0.4 is selected since the optimal value for the correlation coefficient and the RMSE are 0.9997 and 0.2215, respectively.

**Table 4:** Nu-RBF performance utilizing different values of Nu with fixed C=8 and gamma=0.2 of total behaviour prediction model.

| Nu | RMSE |  | Correlation Coefficient |  |  | No. of Support<br>Vectors |
| --- | --- | --- | --- | --- | --- | --- |
|  | Training | Testing | Training | Testing | Overall |  |
| 0.001 | 8.1690 | 8.1908 | 0.8105 | 0.7800 | 0.8026 | 3 |
| 0.1 | 0.5074 | 3.3228 | 0.9984 | 0.9299 | 0.9820 | 63 |
| 0.2 | 0.2386 | 3.2229 | 0.9997 | 0.9346 | 0.9838 | 80 |
| 0.3 | 0.2215 | 3.1969 | 0.9997 | 0.9353 | 0.9840 | 82 |
| **0.4** | **0.2215** | **3.1968** | **0.9997** | **0.9353** | **0.9840** | **82** |
| 0.5 | 0.2219 | 3.1973 | 0.9997 | 0.9353 | 0.9840 | 83 |

There are two important parameters to be selected which are gamma, γ and capacity parameters, C. These are important parameters to optimize the structure of the SVM model to be implemented in certain circumstances. During training stage of the SVM simulation, gamma is fixed to 0.1 and the values of C is set from 0 to 10 (horizontal axis). The error of predictions is evaluated by RMSE and the correlation coefficient along with the number of support vectors (vertical axis). These values are plotted as shown in Fig 7. As the value of C increases, the correlation coefficient also increases while the number of support vectors and RMSE values slightly decrease. In addition, by focusing on the parameter C, the highest value of the correlation coefficient is 0.999. Therefore, the best value of C to be considered at each station is 4.

**Fig 7:**
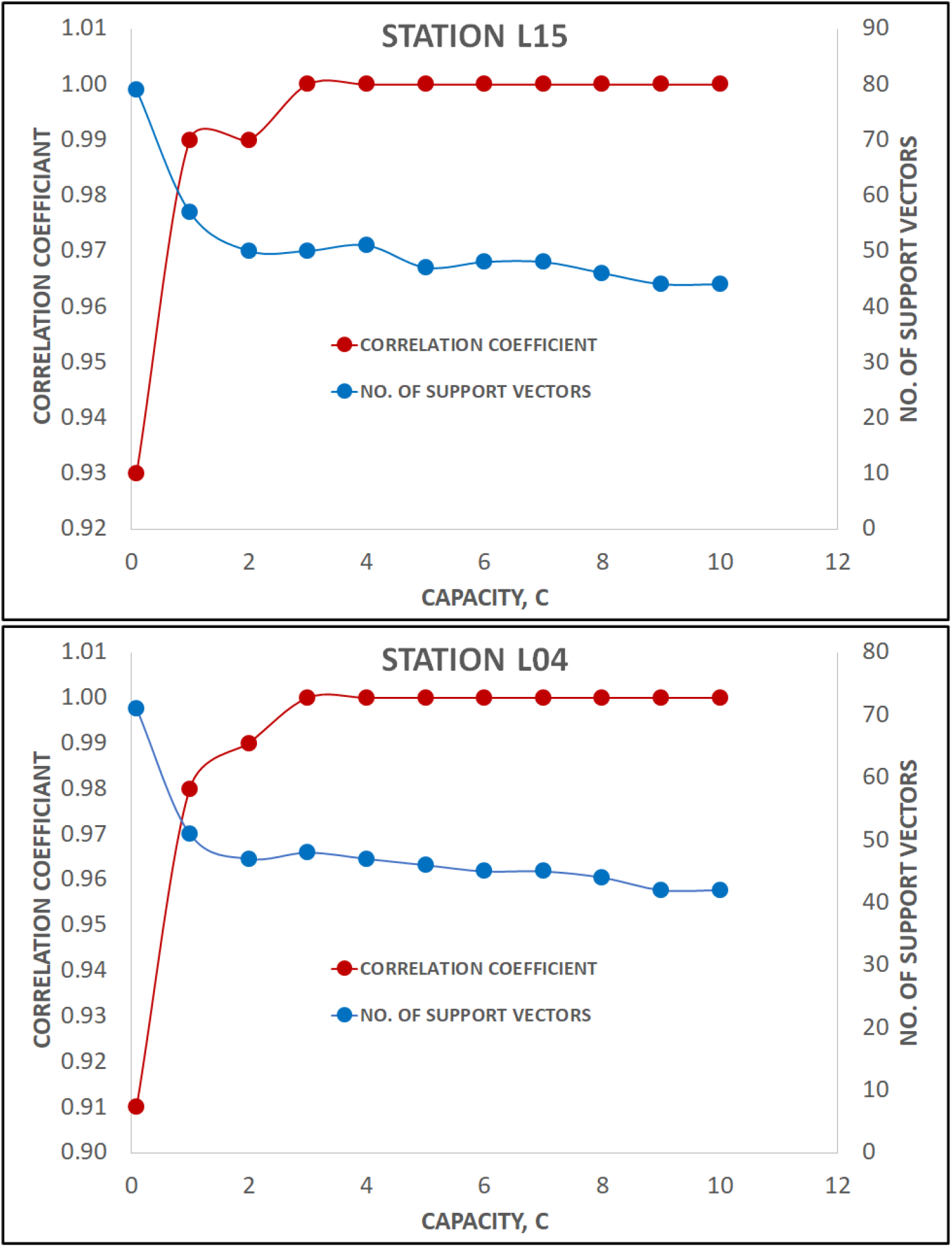
Result of various capacity parameter, C of SVM model for station L15 and station L04.

Generally, it is necessary to determine the optimal values of the hyperparameters C and gamma as a primary step for application of SVM model. Hence, a trial-and-error procedure is needed to estimate the generalization accuracy by utilizing a various values of kernel hyperparameters. Regarding to the limitation of parameter gamma, the optimal values of gamma are decided in the range of 0.001 to 0.9 (vertical axis) with 0.1 unit of gamma increment whereas the values of C=8 and nu=0.4 are fixed. The horizontal axis unchanged as per Fig 7. Initially, 10-fold cross validation is applied to determine the optimal value of gamma with ten repetition to improve reliability of the model results. The final structure of the model is determined as the minimal of RMSE is reached in the testing phase. Fig 8 below shows the relation between correlation coefficient and gamma values as discussed. In the graph, the value of correlation coefficient increases as gamma increase. This consistent trend also followed by the number of support vectors values. However, this trend discontinued for both right and left axis and reach its constant value and remain unchanged at 0.2 for both stations. As a result, the best value to be considered for WQI prediction model in training and forecasting session is when gamma equals to 0.2.

**Fig 8:**
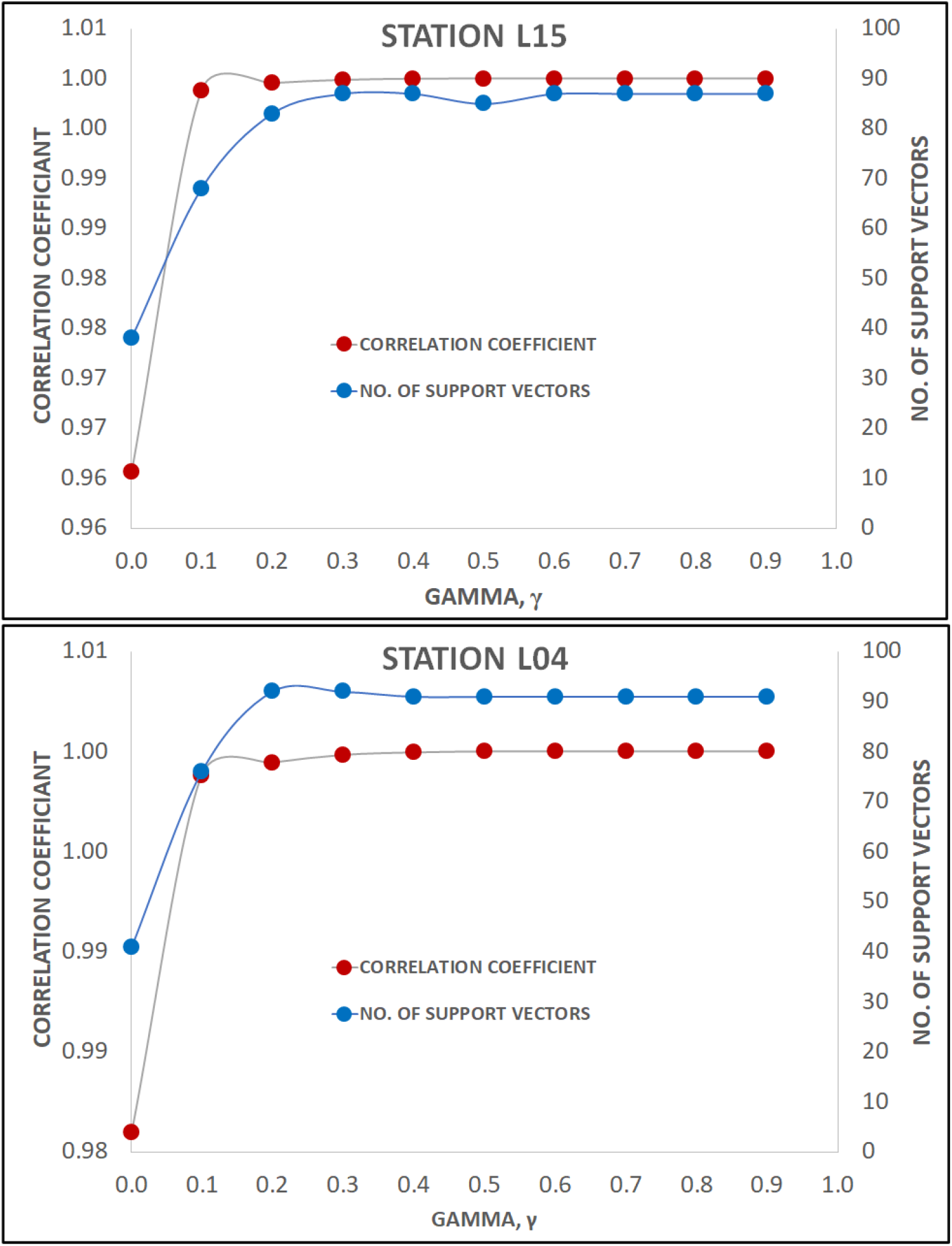
Result of various parameter Gamma, *γ* of SVM model for station L15 and station L04.

K-fold cross-validation is a robust technique specifically designed to evaluate the accuracy of a model. K-fold CV technique functions is to control the test error rate more accurately. In this study, there are a few values of k have been tested through a trial-error procedure. Table 5 below shows statistical evaluation using 3, 5, 7, 10, and 15-fold cross validation for Epsilon-RBF and Nu-RBF models. From the table, RMSE and MAE values of 5-fold cross validation demonstrate the best goodness of fit. The result also indicates the excellent performance for both type of RBF models (R_ε_=0.9994, RMSE_ε_=0.3064, R_Nu_=0.9998, RMSE_Nu_=0.1909). As a comparison, Nu-RBF model produces better accuracy than Epsilon-RBF model in 5-fold CV value. Thus, the Nu-RBF model is selected as the optimal model to evaluate the training data for further analysis.

**Table 5:** Statistical evaluation using 3, 5, 7, 10, and 15-fold cross validation for Epsilon-RBF and Nu-RBF models.

| Statistical Evaluation | k-Fold |  |  |  |  |
| --- | --- | --- | --- | --- | --- |
|  | 3 | 5 | 7 | 10 | 15 |
| ε-RBF Model |  |  |  |  |  |
| RMSE | 0.4054 | **0.3064** | 0.3463 | 0.3907 | 0.3769 |
| MAE | 0.1268 | **0.0804** | 0.1000 | 0.1203 | 0.1141 |
| Correlation Coefficient | 0.9989 | **0.9994** | 0.9992 | 0.9989 | 0.9990 |
| Nu-RBF Model |  |  |  |  |  |
| RMSE | 0.3394 | **0.1909** | 0.4157 | 0.4278 | 0.4105 |
| MAE | 0.0923 | **0.0304** | 0.1818 | 0.2160 | 0.1618 |
| Correlation Coefficient | 0.9992 | **0.9998** | 0.9988 | 0.9988 | 0.9988 |

This selection is supported by a graph as shown in Fig 9. The figure represents the comparison between actual and predicted value of WQI for both stations using the selected Nu-RBF model. The vertical axis represents WQI data and the horizontal axis represents time set. Each plot shows that the previously selected model is capable to predict WQI. This capability is visualized by the similarity behavior of plot between the actual and predicted values of WQI. The closeness of this data indicated that the deviation of error is very small and can be neglected. Hence, this result also confirms the suitability of combination Nu-RBF with 5-fold CV model selection.

**Fig 9:**
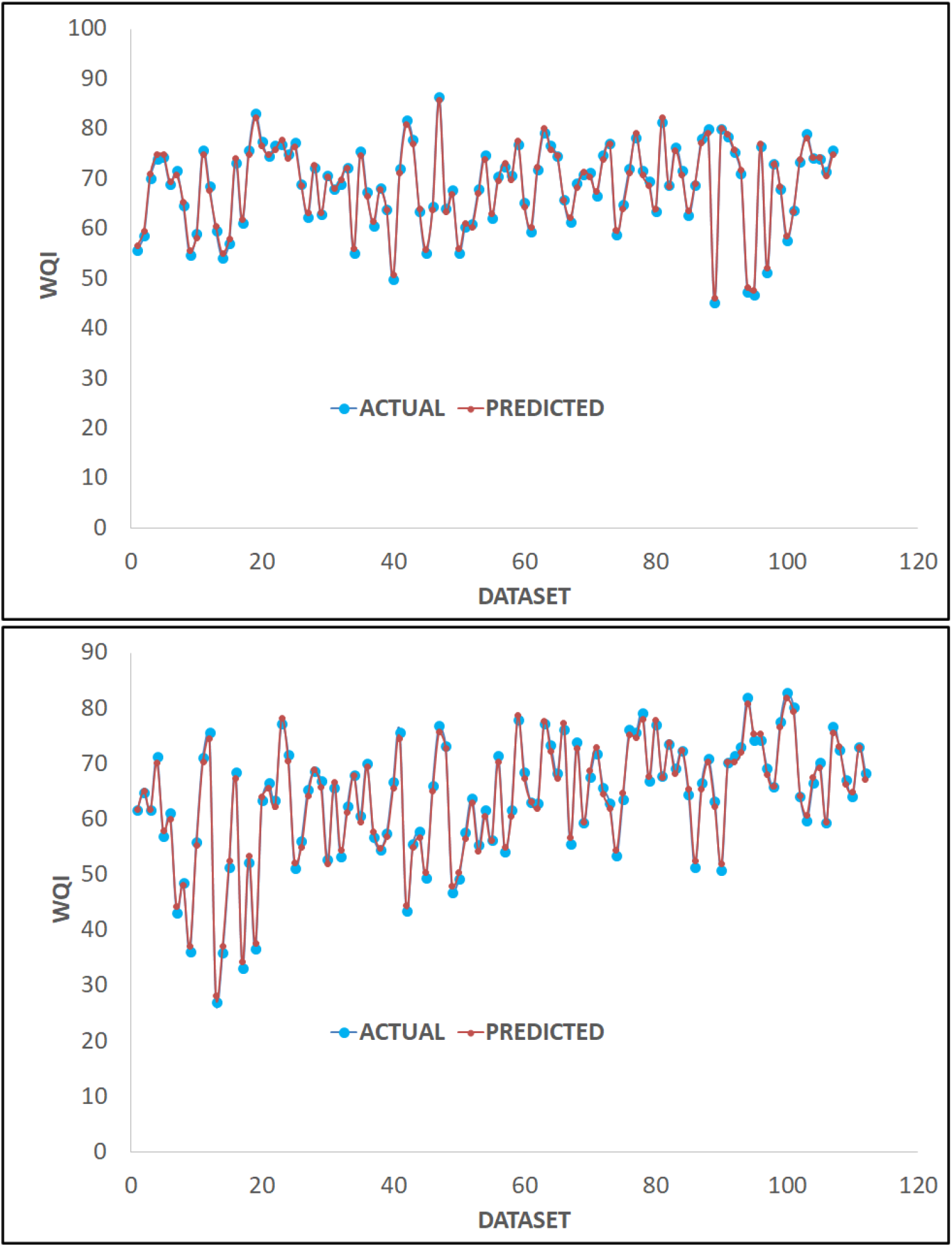
Comparison between observed and predicted WQI for station L15 and station L04.

In Malaysia, WQI is calculated based on DOE-WQI formula by using 6 water quality parameters (ammoniacal nitrogen (AN), biochemical oxygen demand (BOD), chemical oxygen demand (COD), dissolved oxygen (DO), pH, and suspended solids (SS)). If any parameter is missing, the calculation of DOE-WQI is impossible to proceed.

In this study, several input parameter combinations are evaluated to determine the optimal combination for WQI prediction. There are seven possible input parameter combinations (Model 1 - Model 7). The accuracy of the models is evaluated by using the statistical indices such as RMSE, MAE, R and NSE coefficient, for both training and testing phases. Table 6 below shows the performance of the SVR model in training and testing for both station L15 and L04. The performance for Model 1 (station L15: R_Train_=0.961, RMSE_Train_=2.728, R_Test_=0.908, RMSE_Test_=6.398; station L04: R_Train_=0.976, RMSE_Train_=2.516, R_Test_=0.971, RMSE_Test_=6.398) which consist of all 6 parameters can be considered as a benchmark to compare with other model results. As for further stage, all combinations (Model 2 to Model 7) are simulated, and all the results are tabulated in Table 6 below. From the results shown, Model 3 produces the highest prediction performance for both training and testing phases. As can be seen from the table, model 3 is the combinations of DO, BOD, COD, SS and AN where pH parameter is excluded from the list with corresponding results (station L15: RTrain=0.961, RMSE_Train_=2.706, R_Test_=0.916, RMSE_Test_=6.382) and (station L04: RTrain=0.977, RMSE_Train_=2.457, R_Test_=0.979, RMSE_Test_=7.237). Model 3 is chosen because although one parameter is excluded in the analysis, the performance of the model is greater than benchmark model (Model 1) in term of overall indicators.

**Table 6:** Performance evaluations of the SVR model in training and testing phases for station L15 and station L04.

| Phase | Model | Parameters Combination | WQI L15 |  |  |  | WQI L4 |  |  |  |
| --- | --- | --- | --- | --- | --- | --- | --- | --- | --- | --- |
|  |  |  | RMSE | MAE | R | NSE | RMSE | MAE | R | NSE |
| Training | 1 | DO, BOD, COD, SS, pH, AN | 2.728 | 2.159 | 0.961 | 0.893 | 2.516 | 1.865 | 0.976 | 0.940 |
|  | 2 | DO, BOD, COD, SS, pH | 3.829 | 3.219 | 0.927 | 0.790 | 3.353 | 2.480 | 0.956 | 0.894 |
|  | 3 | DO, BOD, COD, SS, AN | **2.706** | **2.140** | **0.961** | **0.895** | **2.457** | **1.795** | **0.977** | **0.943** |
|  | 4 | DO, BOD, COD, pH, AN | 3.630 | 2.978 | 0.928 | 0.811 | 3.334 | 2.560 | 0.958 | 0.895 |
|  | 5 | DO, BOD, SS, pH, AN | 4.017 | 3.470 | 0.931 | 0.768 | 3.193 | 2.557 | 0.966 | 0.904 |
|  | 6 | DO, COD, SS, pH, AN | 3.855 | 2.999 | 0.935 | 0.787 | 3.663 | 2.873 | 0.950 | 0.873 |
|  | 7 | BOD, COD, SS, pH, AN | 3.924 | 3.053 | 0.914 | 0.779 | 4.819 | 3.788 | 0.922 | 0.781 |
| Testing | 1 | DO, BOD, COD, SS, pH, AN | 6.398 | 5.011 | 0.908 | 0.538 | 6.886 | 5.527 | 0.971 | 0.724 |
|  | 2 | DO, BOD, COD, SS, pH | 6.617 | 5.191 | 0.874 | 0.506 | 7.567 | 6.332 | 0.965 | 0.667 |
|  | 3 | DO, BOD, COD, SS, AN | **6.382** | **4.984** | **0.916** | **0.541** | **7.237** | **5.705** | **0.979** | **0.695** |
|  | 4 | DO, BOD, COD, pH, AN | 6.917 | 5.353 | 0.918 | 0.461 | 6.878 | **5.523** | 0.972 | 0.725 |
|  | 5 | DO, BOD, SS, pH, AN | 7.440 | 5.887 | 0.912 | 0.376 | 8.496 | 6.452 | 0.947 | 0.580 |
|  | 6 | DO, COD, SS, pH, AN | 7.229 | 5.585 | 0.925 | 0.411 | 8.628 | 6.767 | 0.965 | 0.567 |
|  | 7 | BOD, COD, SS, pH, AN | 6.923 | 5.387 | 0.858 | 0.460 | 8.876 | 7.318 | 0.936 | 0.542 |

Meanwhile, if DO is omitted from the analysis, the result of WQI shows the lowest accuracy. This lowest accuracy is shown by Model 7 results (station L15: R_Train_=0.914, RMSE_Train_=3.924, R_Test_=0.922, RMSE_Test_=4.819; station L04: R_Train_=0.858, RMSE_Train_=6.923, R_Test_=0.936, RMSE_Test_=8.876). This finding is very significant because it shows that DO cannot be excluded from the analysis because the overall WQI performance will be jeopardize.

The prediction performance of Model 3 for station L15 and station L04 have been plotted in Fig 10. The vertical axis represents predicted WQI and the horizontal axis represents actual WQI. The scatter plot graphically illustrates the association between the predicted and the observed values of WQI. In the diagram, the line of the best fit is drawn as a reference and to describe the closeness of the relationship between the predicted and the observed data. As expected, Model 3 shows excellent prediction performance as the data points is very close to the best fit line in both (training and testing) stations.

**Fig 10:**
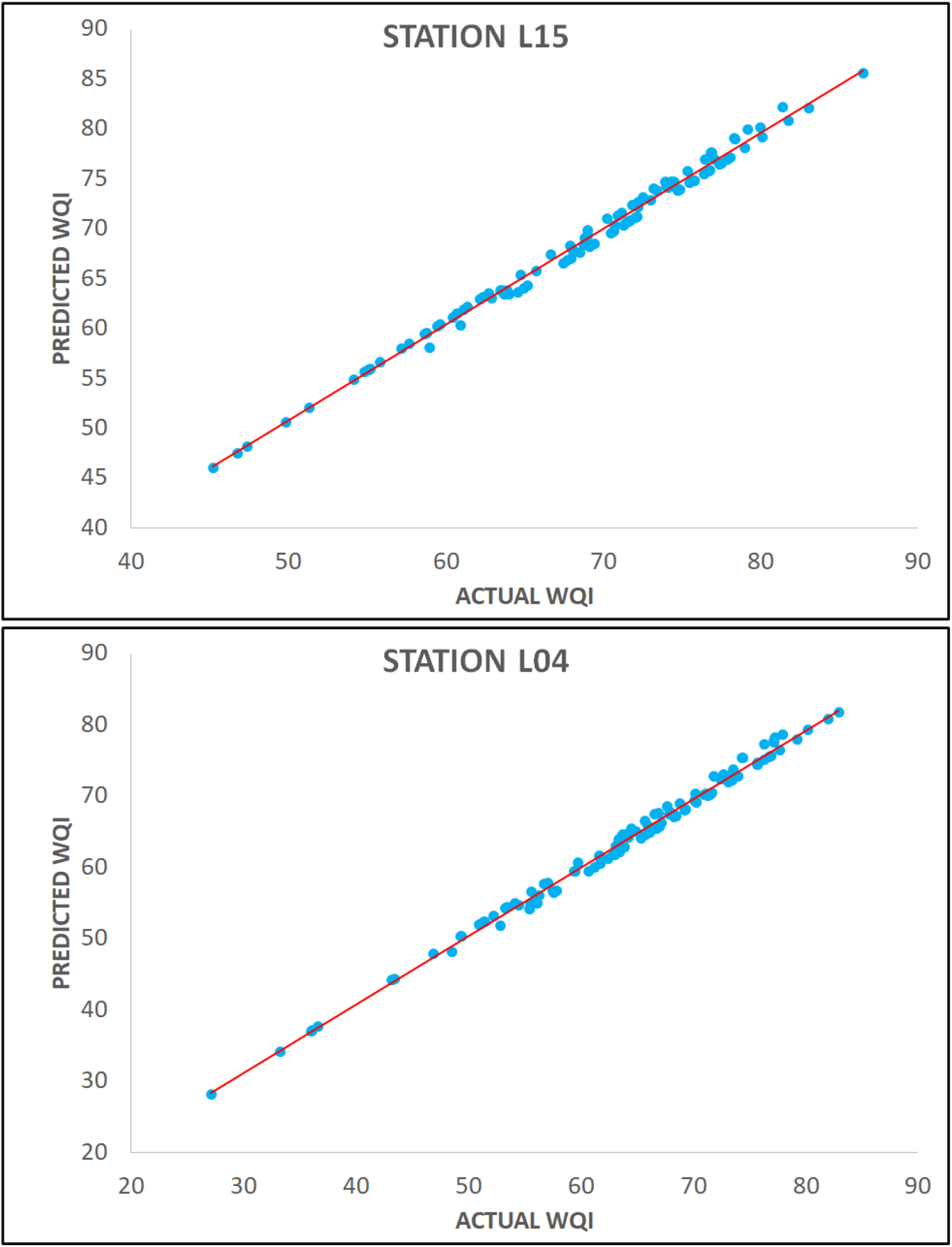
Prediction performance of WQI by using SVR-Model 3 for station L15 and station L04.

## Conclusion

The prediction of water quality is very crucial in pollution monitoring and to ensure the environmental standards for water resources are being met. If there are changes in water quality, this prediction can provide early warnings and thus possibly minimize the effects of poor water quality to the community if stringent action is executed properly by the authority. In this study, two-stations monthly water quality data from Langat River, Kajang are utilized by using six water quality parameters for ten-year period.

In this research, the Support Vector Regression (SVR) model combined with K-fold cross-validation is proposed to predict the Water Quality Index (WQI). Table 5 previously shows the Nu-RBF type with 5-fold CV provides the highest correlation coefficient (R), 0.9998 compared to Epsilon-RBF, 0.9994. This result concludes that the optimal performance of SVR selection is obtained through 5-fold cross-validation and this value of K-fold CV provides better performance over the other K-fold CV value.

The SVR algorithm is developed by considering several input parameters combination (Model 1 – Model 7), whereby each model used a different combination of water quality parameters as modelling input (Table 6). As per the analysis, the best prediction accuracy is achieved by the SVR Model 3 for both stations whereas the R-values are 0.961 and 0.977, respectively. Practically, when one parameter can be omitted from WQI prediction, the whole process can be reduced to an effective cost and time. Meanwhile, the SVR Model 7 (neglecting of DO parameter) shows the lowest accuracy in all performance indicators (Table 6). This scenario proved that the most important parameter that affects WQI is DO and the least important variable is pH. This is another significant finding whereby DO is excluded from the analysis then the overall WQI performance will deteriorate.

As for concluding remarks, the hybrid of Support Vector Regression Model with 5-fold cross-validation is an accurate tool for WQI prediction at Langat River, Kajang. This statement is proved by the accuracy performance values of Model 3 for both stations in training and testing phases. As a result, the RMSE and MAE prediction data can be minimized and consequently the overall SVR prediction can be optimized.

## Acknowledgments

The authors would like to thank the Earth Observation Centre, Universiti Kebangsaan Malaysia, and the Department of Environmental Malaysia (DOE), Ministry of Natural Resources and Environment, Malaysia for providing the data for this research.

